# Opening the side exit pores of ClpP by lowering the pH of proteolytic chamber coupled with substrate hydrolysis

**DOI:** 10.1101/2021.09.20.461017

**Authors:** Leehyeon Kim, Byung-Gil Lee, Min Kyung Kim, Do Hoon Kwon, Hyunmin Kim, Heike Brötz-Oesterhelt, Soung-Hun Roh, Hyun Kyu Song

**Affiliations:** Department of Life Sciences, Korea University, Seoul 02841, South Korea; School of Biological Sciences, Institute of Molecular Biology and Genetics, Seoul National University, Seoul 08826, South Korea; Interfaculty Institute of Microbiology and Infection Medicine, Department of Microbial Bioactive Compounds, University of Tuebingen, Tuebingen, Germany; Cluster of Excellence Controlling Microbes to Fight Infection, University of Tuebingen, Tuebingen, Germany

**Keywords:** acyldepsipeptide, asymmetric binding, ClpP, conformational change, cryo-EM, crystal, pH drop, protein degradation, proteolytic chamber

## Abstract

The ClpP serine peptidase is a tetradecameric degradation machine involved in many physiological processes. It becomes a competent ATP-dependent protease with Clp-ATPases. Small chemical compounds, acyldepsipeptides (ADEPs), are known to cause dysregulation and activation of ClpP without ATPases, and have potential as novel antibiotics. Previously, structural studies of ClpP from various species revealed the structural details, conformational changes, and activation mechanism. Although product release by the side exit pores has been proposed, the detailed driving force for product release remains elusive. Here, we report crystal structures of ClpP from *Bacillus subtilis* (BsClpP) in unforeseen ADEP-bound states. Cryo-electron microscopy structures revealed various conformational states at different pH conditions. To understand the conformational change for product release, we investigated the relationship between substrate hydrolysis and the pH lowering process. Our data, together with previous findings, provide insight into the molecular mechanism of product release by ClpP self-compartmentalizing protease.

## Main

Energy-dependent proteases are molecular machines conserved in all kingdoms of life^1,2^. They play an essential role in protein quality and quantity control by degrading misfolded, damaged, short-lived or regulatory proteins using energy by ATP hydrolysis^3,4^. Clp proteases are an energy-dependent protease system in bacteria and mitochondria of eukaryotic cells^5^. They contain two distinct functional components, a Clp-ATPase and the ClpP proteolytic core. The energy-consuming hexameric AAA+ ATPase (ATPase associated with a variety of cellular activities) is responsible for substrate selection, unfolding, and translocation into the proteolytic core. Representatives include ClpA, ClpC, ClpE, and ClpX. The central proteolytic machine ClpP is a barrel-like serine protease complex, composed of two stacked heptameric rings that form an enclosed degradation chamber^6^. Both ClpP heptamers have a central axial pore where the unfolded polypeptide chains are translocated.

The structures of ClpP from various species, from *E. coli* to humans, have been solved, and information regarding folding, the axial entrance pore, and the catalytic triad of ClpP is well defined^6–13^. The activation mechanism of ClpP by its Clp-ATPase partner was a long-standing question in the field and partially revealed by structures of ClpP in complex with acyldepsipeptides (ADEPs)^11,14^, compounds that have antibiotic activity through dysregulation of ClpP activity by mimicking the Clp-ATPase^15–17^. Very recently, more direct evidence comes from the complex structures formed between the ClpX ATPase and ClpP protease using cryo-electron microscopy (cryo-EM)^18–20^. These studies showed how the symmetry-mismatched ATPase activates the proteolytic machine by translocating substrate molecules. The critical IGF/L (Ile-Gly-Phe in ClpX and Ile-Gly-Leu in ClpA) loops of the Clp-ATPases bind to the hydrophobic pockets between the ClpP monomers, as shown in the ADEP-bound structures. The asymmetric ClpAP complexes showed one or two empty IGL-loop binding pockets during the engagement and disengagement cycle^21^, which had not been clear in the previously determined structures of ClpP fully occupied with ADEP compounds^11,14,22^. The activation mechanism of ClpP by its cognate ATPases has now been proposed. However, the detailed and coherent steps of ClpP motion, from substrate entry to product release, are still elusive, although several conformational equilibria between active and inactive states have been reported, such as pH-dependent switching, active site perturbation, and regulation of the N-terminal loop^23–26^.

Once substrates reach the tetradecameric proteolytic chamber of ClpP, the degraded products need to be eliminated for efficient processing. The equatorial pore for peptide release was structurally revealed by the observation of major conformational changes of ClpP within the same strains ^13,25,27,28^, corroborated by an elegant biophysical study using NMR. Currently, several other factors have been proposed for the regulation of equatorial pore opening ^23–26,30^. However, those studies could not fully explain the peptide product release mechanism of ClpP coupled with its activators, AAA+ ATPases or ADEPs. Here, we present the crystal and cryo-EM structures of ADEP-bound ClpP in various states and a plausible link between a decrease in pH and peptide hydrolysis, which reveal conformational changes and the peptide release mechanism of ClpP. These data, in combination with all available structural information, show the complete processing of substrates, from recognition to unfolding, translocation, degradation, and release.

## Results

### Lowering the pH by the accumulation of hydrolyzed peptide products

As described, there are significant conformational changes in the handle region of ClpP from *Bacillus subtilis* (BsClpP), as shown in our previous studies^11,27^, and those from other species ^22–24,26,28^. It has been proposed that His145 interacting with the catalytic aspartate of an adjacent protomer in *N. meningitidis* ClpP serves as a key determinant for a pH-dependent conformational switch^23^. However, the equivalent residue in BsClpP is alanine, and therefore, there must be differences among species (Extended Data Fig. 1). In *M. tuberculosis* ClpP1P2, peptide binding at the active site triggers the transition between active and inactive conformations^24^. Therefore, it is tempting to speculate that there must be a relationship between pH change and peptide accumulation in the proteolytic chamber of ClpP. Furthermore, there have been reports that the pH drop can be correlated with peptide bond hydrolysis^31,32^, and naturally, the numerous newly generated amino- and carboxy-terminal groups contribute to the pH drop from neutral pH values. The average pI values of bacterial proteomes are also slightly acidic^33^. Therefore, we hypothesized that peptide bond hydrolysis decreases the pH value of a protein solution. To test this hypothesis, we checked the pH change during protein hydrolysis. The initial pH of 6.41 achieved using diluted PBS (phosphate-buffered saline) with 80 μM α-casein decreased to the terminal pH of 5.94 by the addition of proteinase K. The drop of 0.47 pH units occurred very fast because of the high enzymatic activity of proteinase K for the partially unfolded α-casein substrate (Fig. 1a and Extended Data Fig. 2a). We observed a similar pH change with BsClpP protease in the presence of ADEP1 (Fig. 1a). In contrast to monomeric proteinase K, the proteolytic activity of BsClpP is lower, and thus, the pH drop was slower. Nonetheless, a substantial drop by 0.46 pH units was reached from the initial pH 6.58 to pH 6.12 after 300 min (Fig. 1a). The same experiment performed with bovine serum albumin (BSA) showed an even stronger pH drop of 0.76, from 6.80 to 6.04 (Fig. 1b). In the absence of a protease, the pH values were quite stable (Fig. 1). In parallel to the pH measurement, protein degradation was followed by SDS-PAGE (Extended Data Fig. 2). The pIs of intact α-casein and BSA are 4.91 and 5.32, respectively. These values are derived from the summation of pK_a_ values of all side-chain groups and amino- and carboxy-terminal groups. However, when proteases cleave numerous peptide bonds in the proteins, many new α-amino and carboxylic groups are generated. Subsequently, the pH value of the environment is mainly governed by the total summation of all pK_a_ values of α-amino and carboxylic groups and is below 6.0, which is slightly acidic. The cleaved peptide products must be highly concentrated in the proteolytic chamber of ClpP, and therefore, the local pH of the handle region near the active site becomes acidic. Consequently, we conclude that the conformation of BsClpP at low-pH conditions mimics the state during peptide accumulation inside the proteolytic chamber.

**Fig. 1:**
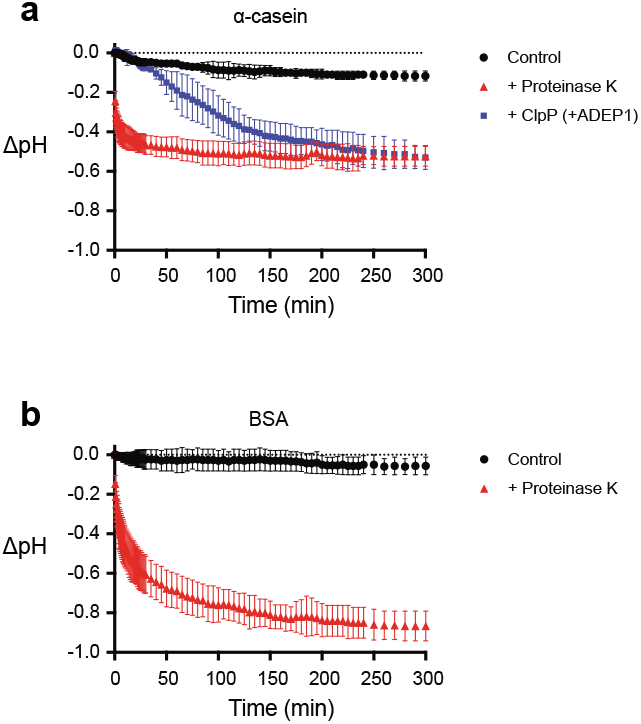
pH change during protein hydrolysis. **a,** Monitoring of the pH change during degradation of α-casein by BsClpP in the presence of ADEP1 and proteinase K. Blue (square) and red (triangle) symbols represent α-casein with BsClpP+ADEP1 and proteinase K, respectively. **b,** Monitoring of the pH change during degradation of bovine serum albumin (BSA) by proteinase K (red triangle). Black circles represent the control, only substrates without proteases (**a,b**). Data points represent the mean value of three measurements and the error bars show standard deviations.

### Asymmetric ADEP binding in crystal structures of the BsClpP-ADEP complex

We solved two structures of BsClpP in complex with acyldepsipeptides (specifically ADEP2) at 2.8 and 3.0 Å resolution (Fig. 2 and Supplementary Table 1). For molecular replacement, previously compressed BsClpP (PDB ID: 3TT6) was used as a search model^27^. Very intriguingly, these two structures have two and five ADEP2 compounds, respectively, bound to each heptameric ring (Fig. 2 and Extended Data Fig. 3), which is in stark contrast to the previous 14 ADEP-bound BsClpP^11^ as well as all other ADEP-ClpP complex structures of from various organisms published to date^22,34–36^. Our two new BsClpP tetradecamer structures have a large difference in their degrees of compression as well as in their numbers of ADEP molecules bound, four and ten (Fig. 2). Hereafter, 2ADEP and 5ADEP indicate 2 and 5 ADEPs for each heptameric ring (4 and 10 ADEPs for a BsClpP tetradecamer). Their axial heights are approximately 86 Å (2 ADEPs) and 92 Å (5 ADEPs), and the state of BsClpP bound to 4 ADEPs (2ADEP) has a similar height to the previous compressed apo-form BsClpP structure^27^. For consistency, the height was defined as the distance between the outermost two atoms; therefore, it is slightly taller than that in previous reports, which is usually the distance measured between the two most distant Cα atoms. The height of BsClpP in complex with 10 ADEPs (5ADEP) is in the middle of the extended and compressed states, which is similar to the state called the ‘compact state’ ^13,35,37^. Thus, these two different states are referred to as ‘compressed’ for the 2ADEP structure and ‘compact’ for the 5ADEP structure. In both structures, ADEPs bind at the interface of two subunits in the order of A-E-E-E-A-E-E (A: ADEP-bound and E: empty) in the compact state and A-A-A-E-A-A-E in the compressed state (Fig. 2).

**Fig. 2:**
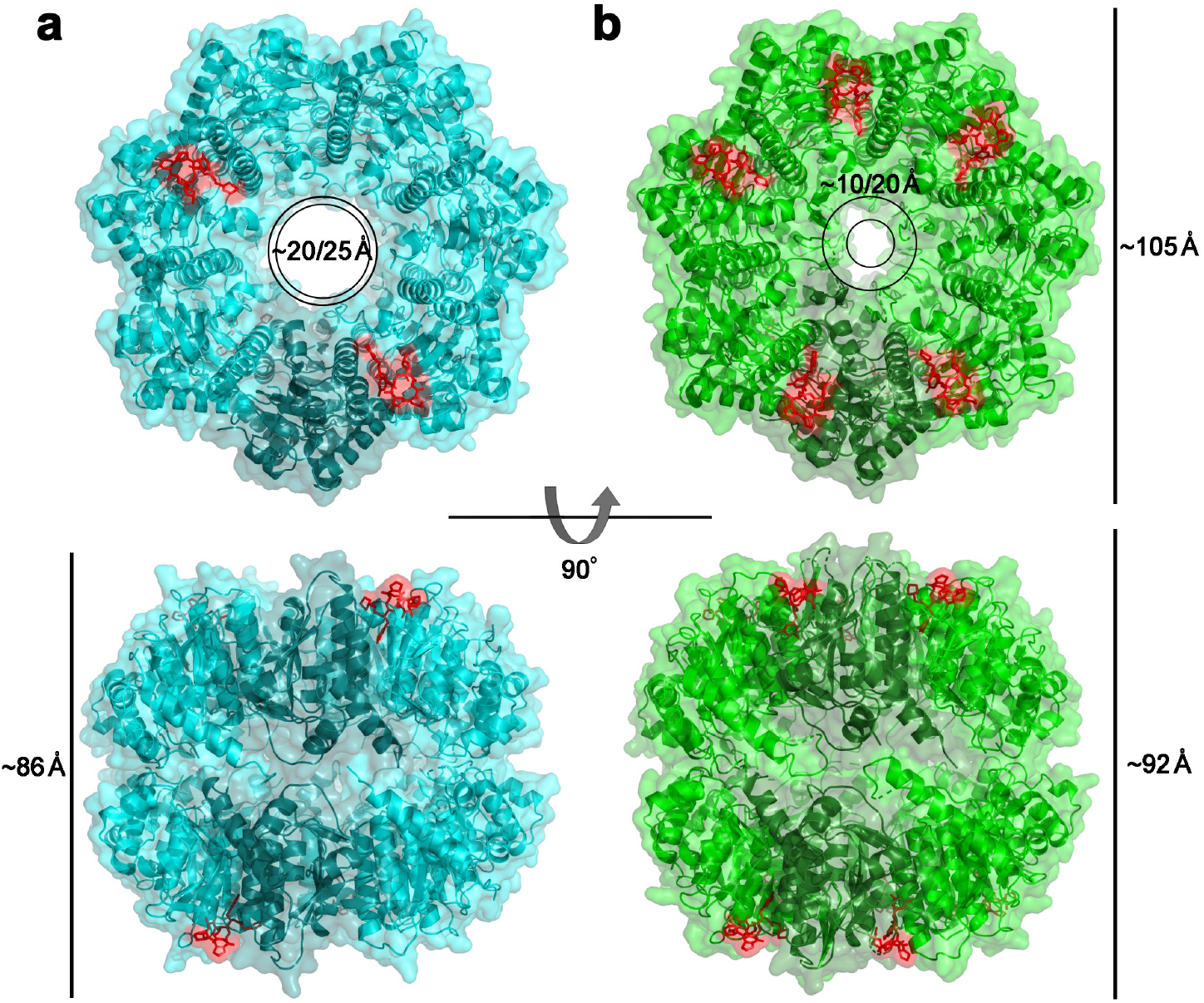
X-ray structures of ADEP-bound BsClpP. **a,** Ribbon diagram of 2ADEP-bound BsClpP (cyan) with a transparent molecular surface and one monomer in each heptameric ring, colored darker for clarity (2ADEP). **b,** Ribbon diagram of 5ADEP-bound BsClpP (green) with a transparent molecular surface and one monomer in each heptameric ring, colored darker for clarity (5ADEP). Tetradecameric BsClpP is viewed along a 7-fold molecular symmetry axis (upper), and the 2-fold side view is observed by rotating 90° (lower). The bound ADEP molecules colored red are shown as a stick model. Dimensions of the models are indicated. The two diameters are noted due to the asymmetric shape of the entrance pore.

### Different compressions of BsClpP control the entrance and exit pores

These two states differ in terms of the preservation of N-terminal residues in addition to height. The N-terminal segments of compact 5ADEP are well structured and resolved, whereas those of 2ADEP are disordered, so the pore sizes in the two states are markedly different (Fig. 2). Due to the asymmetric binding of the activator ADEP, the entrance pore showed distorted shapes in both states, and in particular, the compact state possessed even more asymmetry, in that the diameter between empty subunits was nearly half that of ADEP-bound subunits (Fig. 2b). More importantly, the pores in the lateral direction, known as the product exit, were also different. This correlates with the flexibility or unwinding of the α5-helix, called the handle region ^13,27,28^. Since we obtained asymmetric ADEP-bound structures, each monomer in different neighboring subunits was compared (Fig. 3 and Extended Data Fig. 4). In both structures, each subunit has three different environments, and details of the 2ADEP and 5ADEP structures are different. In the orientation looking down from the substrate entrance pore in the case of 2ADEP (Fig. 3a), there are three states: protomers having ADEP on the right side (2 subunits), ADEP on the left side (2 subunits), and no ADEP binding (3 subunits). The monomeric structures of protomers with ADEP bound at either the right or the left sides are quite similar, whereas the unbound monomers are slightly different, especially regarding the loop right before the handle region (Fig. 3b). The N-terminal regions of all subunits are quite flexible and have an almost invisible electron density; thus, the asymmetry of the entrance pore is only marginal (Fig. 2a).

**Fig. 3:**
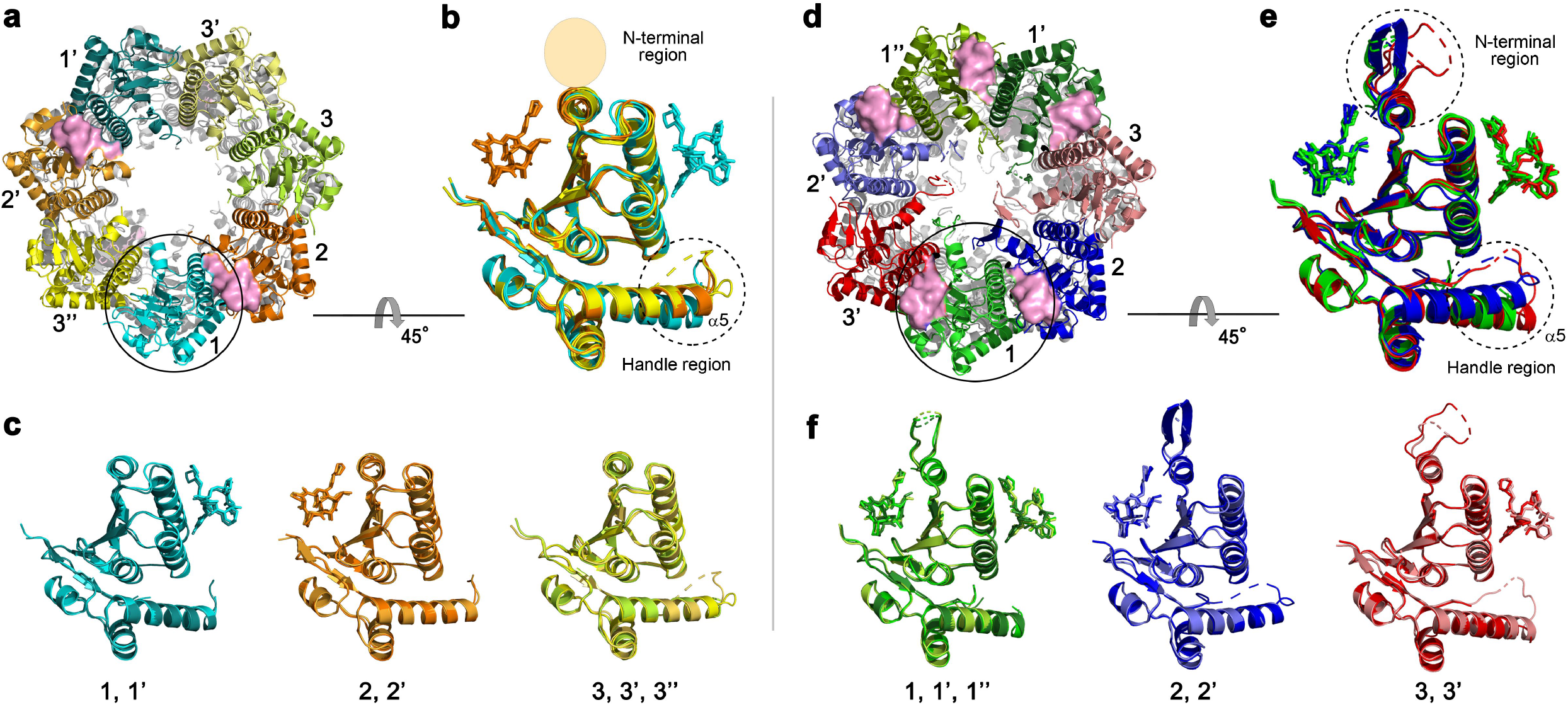
Structural analyses of the compressed 2ADEP and the compact 5ADEP. **a,** Three different subunit environments of 2ADEP viewed along a 7-fold axis: two cyanish subunits (1 and 1’) with the bound ADEP (pink molecular surface) on the right side, two orangish subunits (2 and 2’) with the bound ADEP on the left side, and three yellowish subunits (3, 3’ and 3’’) with no ADEP molecule. **b,** Superposition of all 7 subunits in the heptameric ring of 2ADEP viewed by rotating the panel (**a**) 45° about the horizontal axis. The invisible N-terminal region, due to flexibility, is marked with a transparent oval, and the structurally dynamic handle region is marked with a dashed circle. **c,** Superposition of cyanish subunits (1, 1’), orangish subunits (2, 2’), and yellowish subunits (3, 3’, 3’’). The view is the same as that of panel (**b**). **d,** Three different subunit environments of 5ADEP viewed along a 7-fold axis: three greenish subunits (1, 1’ and 1’’) with the bound ADEP (pink molecular surface) on both the left and right sides, two bluish subunits (2 and 2’) with the bound ADEP on the left side, and two reddish subunits (3 and 3’) with ADEP on the right side. **e,** Superposition of all 7 subunits in the heptameric ring of 5ADEP viewed by rotating the panel (**d**) 45° about the horizontal axis. The very flexible N-terminal region and handle region are marked with dashed circles. **f,** Superposition of greenish subunits (1, 1’, 1’’), bluish subunits (2, 2’), and reddish subunits (3, 3’). The view is the same as that of panel (**e**).

In the same orientation of 5ADEP (Fig. 3d), there are also three different states: ADEP on both sides (3 subunits), ADEP on the right side (2 subunits), and ADEP on the left side (2 subunits). Therefore, the main difference between 2ADEP and 5ADEP is that 2ADEP has 3 monomeric subunits with ADEP on neither side and 5ADEP has 3 monomeric subunits with ADEPs on both sides. In the case of 5ADEP, the monomeric structures with ADEP bound on both sides are quite similar (Fig. 3f), whereas the monomers with a single ADEP on either the right or the left side are different in the loop right before the handle region and in the N-terminal regions (Fig. 3e). The superposition of all subunits in 5ADEP shows that the N-terminal regions and the α5 handle region of all subunits are very dynamic, and thus, the asymmetry of the entrance pore, as well as the side pores, is augmented (Figs. 2b, 3e, and Extended Data Fig. 4).

### Diverse ADEP-bound states found in cryo-EM structures

Although the 2ADEP and 5ADEP states in crystals must be intermediate structures in the reaction cycle of ClpP, we performed cryo-EM experiments at the pH of the crystallization buffer to rule out the possibility of crystal packing artifacts. We obtained 2ADEP and 5ADEP structures with pH values of 4.2 and 5.6, respectively. Unfortunately, at pH 5.6, BsClpP formed an aggregate, probably due to its pI of 5.2; therefore, we collected cryo-EM data at pH 6.5 and 4.2. From datasets of the BsClpP particles, we were able to reconstruct 4 different structures of BsClpP at 3.1, 3.2, 3.4, and 3.4 Å maps (Extended Data Fig. 5 and Supplementary Table 2). The initial electron micrographs showed heterogeneous ADEP-bound states. The 1:1 mixture between BsClpP and ADEP1 at a pH of 6.5 showed that the majority of the particles were apo forms (Fig. 4a and Extended Data Fig. 5a), whereas in the 1:3 mixture (i.e., three-fold surplus of ADEP1) at the same pH, the majority were 14 ADEP-bound forms (Fig. 4b and Extended Data Fig. 5b). The apo structure at pH 6.5 is similar to that of the previous extended structure, and the ADEP-bound structure showed a very clear N-terminal region (Extended Data Fig. 6). In the electron micrographs of the 1:3 mixture of BsClpP and ADEP at pH 4.2, both apo and all ADEP-bound BsClpPs were shown as significant populations (Extended Data Fig. 5c); thus, we determined the two structures with one dataset (Fig. 4c,d, and Supplementary Table 2). However, as shown in Extended Data Fig. 5, other complex states also existed, and we believe that the 2ADEP and 5ADEP found in crystals are intermediates that are energetically stable in the conformational transition from extended to compressed.

**Fig 4:**
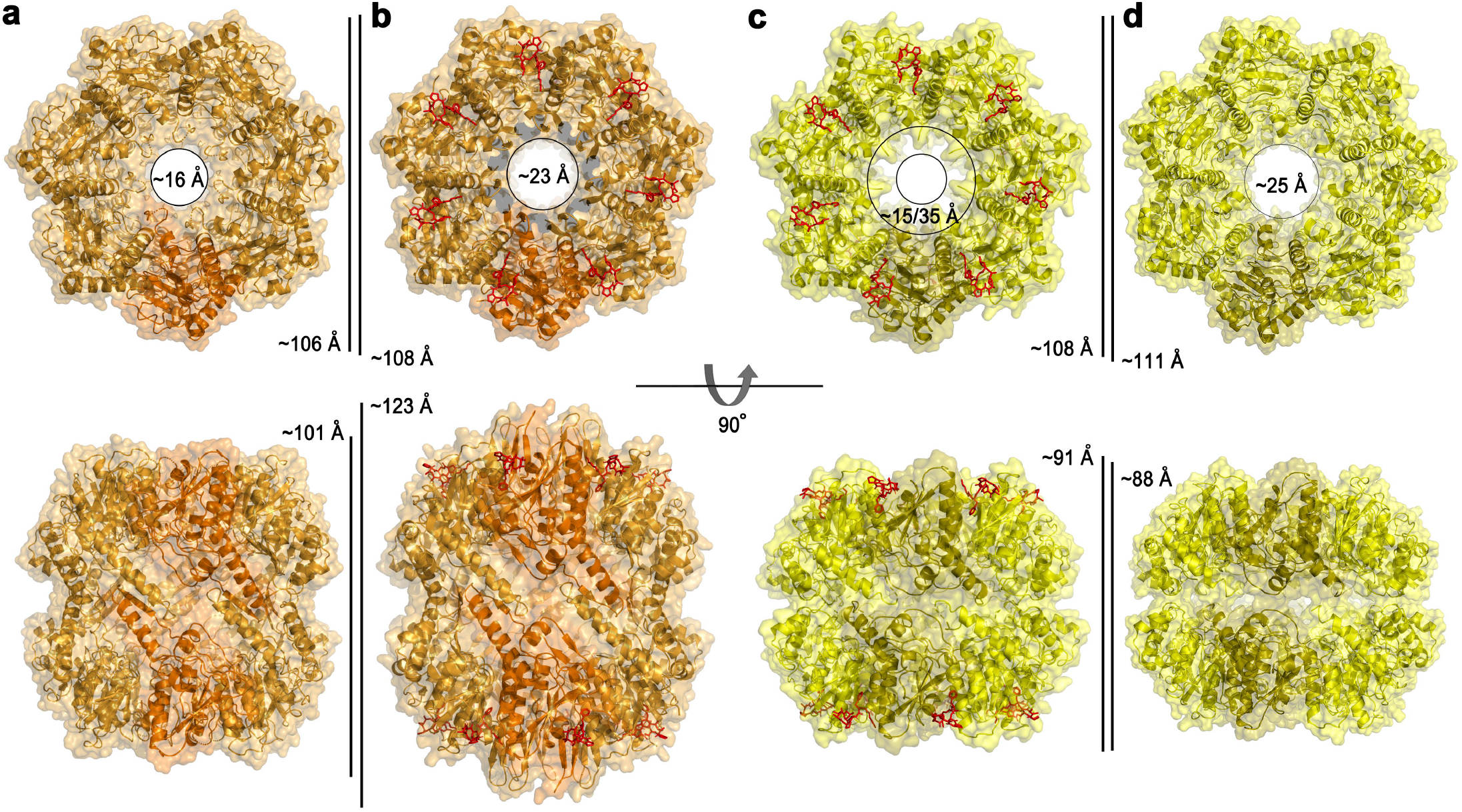
Cryo-EM structures of the BsClpP-ADEP1 complex. **a,** Ribbon diagram, with the transparent molecular surface of the extended apo-BsClpP structure at pH 6.5 viewed along a 7-fold molecular symmetry axis (upper row) and viewed with 90° rotation to display the side view of the 2-fold symmetry axis (lower row). One monomer is shown in darker orange color for clarity. **b,** Same representation as that of panel (**a**) showing the 14 ADEP1-bound BsClpP at pH 6.5. The bound ADEP1 molecules are shown as stick models colored red. **c,** Same representation of the compressed BsClpP with 14 ADEP1 molecules at pH 4.2. One monomer is shown in darker yellow color for clarity. **d,** Same representation of the compressed apo BsClpP at pH 4.2. Dimensions of the models are indicated.

The 14 ADEP-bound BsClpP cryo-EM structures at the two different pH values, 6.5 and 4.2, are markedly different in dimensions (Fig. 4b,c). The structure at pH 6.5 shows an extended conformation, with a well-ordered N-terminal region slightly tilted to the outward position, whereas the 14 ADEP-bound structure at pH 4.2 is similar to the compact structure, with a relatively ordered N-terminal region to the inward position, suggesting that the N-terminal region might be involved in the substrate feeding process. The N-terminal region of ClpP acts as a gate for controlling substrate access with Clp-ATPases as well as ADEP activators ^26,35,38,39^. This ADEP-bound structure at pH 4.2 is classified as a compact conformation based on its dimension, and it is very similar to 5ADEP (Fig. 2b). However, the entrance pore is symmetric because all subunits are virtually identical (Fig. 4c). The apo BsClpP structure at pH 4.2 is structurally similar to 2ADEP (Fig. 2a), although details, such as the entrance pore and handle regions, are subtly different.

### Structural changes in the proteolytic chamber

To understand more details about the various structural states at different pH conditions, we focused on the interior of the proteolytic chamber. Although key determinants for the maintenance of the active extended conformation of ClpP, including the β-sheet in the handle regions of two subunits from each heptameric ring and critical salt-bridge pairs (Asp169– Arg170’:Asp169’–Arg170) at the tip of handle region, have been extensively studied, the charge property of the chamber has not been examined thoroughly. We analyzed the electrostatic potential surface of all different structural states from the perspective of hydrolyzed substrates (Fig. 5). In general, the surface shows negatively charged features in all states, and it can be an environment in which the negatively charged products are energetically unfavorable, although they are just slightly acidic due to the summation of the pK_1_ (–COOH), pK_2_ (–NH_3_^+^), and pK_R_ (–R: side chain) of all peptide fragments, as we described in an earlier section (Lowering the pH by the accumulation of hydrolyzed peptide products). To release this strained state derived from product generation, a conformational change must be triggered. Usually, the side chain of the histidine residue (pK_R_ = 6.0) is thought to be a sensor for the pH-dependent transition. Previously, His145 in *N. meningitidis* ClpP was proposed to be a switch regulating the pH-dependent conformational change and it forms an inter-subunit H-bond with the catalytic Asp178’ residue from a neighboring ClpP protomer^23^. However, the equivalent residues in ClpPs from *B. subtilis, M. tuberculosis,* and *S. aureus* are alanine and glutamine, which are not conserved (Extended Data Fig. 1). Assuming the conservation of the molecular motion of ClpP among all species, the key switch residues must be strictly conserved. Therefore, we thought that another histidine residue, and particularly the strictly conserved catalytic His122, might be a candidate (Extended Data Fig. 1). The catalytic triad Ser97-His122-Asp171 is an active configuration in the extended conformation (Fig. 5a,b), whereas the compact and compressed conformation possesses an inactive configuration (Fig. 5c-f). Naturally, the imidazole ring of His122 is protonated at low pH, and subsequently, the catalytic triad is severely distorted. In the compact and compressed structure, the catalytic triads in the upper and lower heptameric rings close, and the previously identified key residues do not participate in the maintenance of the extended structure. The generation of exit side pores is coupled with the transition from an extended state to a compact state (Fig. 5c,d), and the pores for peptide release are gradually enlarged to a compressed state (Fig. 5e,f).

**Fig. 5:**
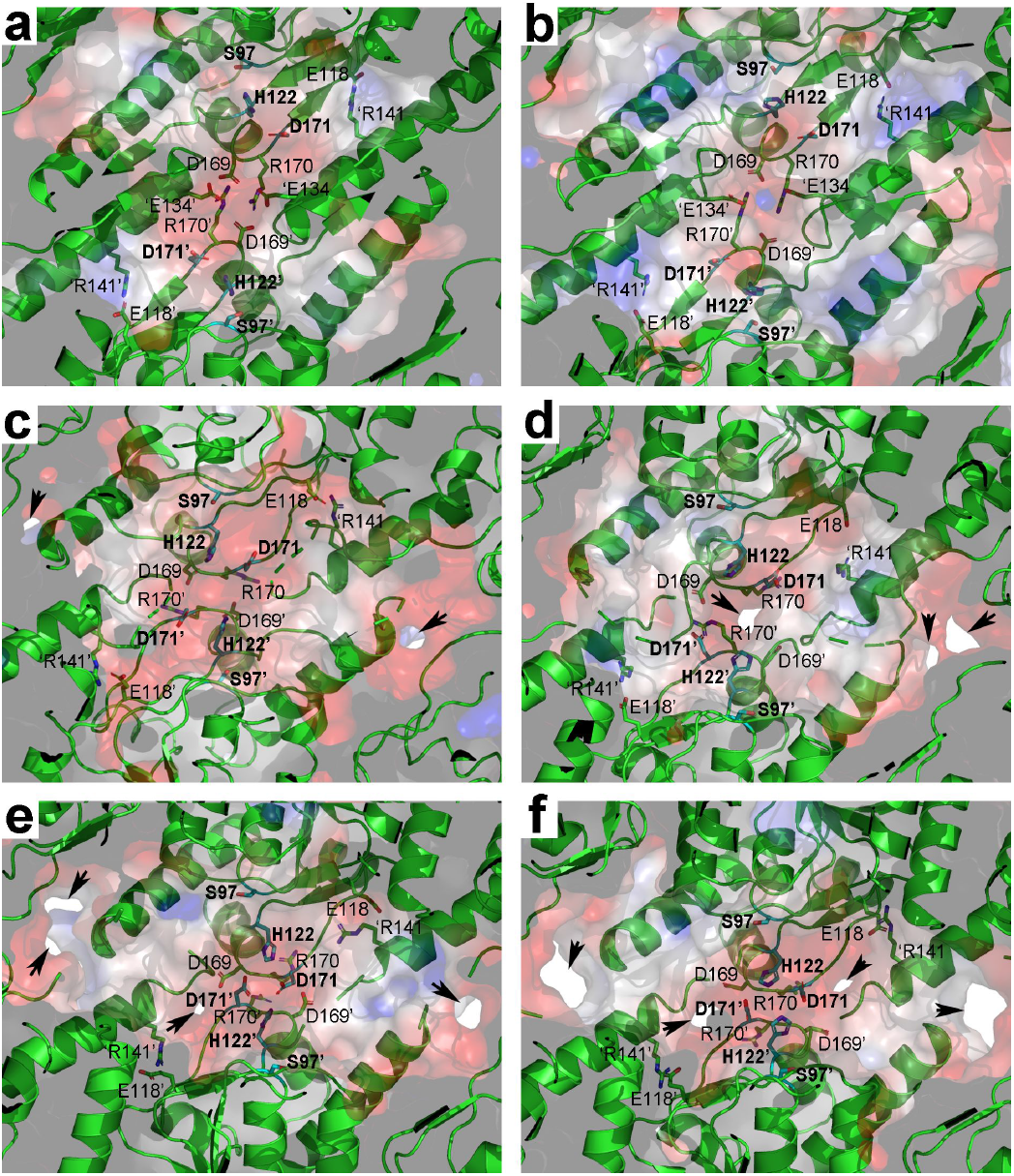
Electrostatic potential of the proteolytic chamber of various structural states. **a,** Ribbon diagram, with the transparent electrostatic potential surface of extended BsClpP viewed from the center of the proteolytic chamber. The catalytic triad (S97-H122-D171) shows the fully active configuration. **b,** Same representation of the 14 ADEP-bound BsClpP cryo-EM structure at pH 6.5. The residues maintaining the extended conformation are now slightly loosened, and the catalytic triad is slightly distorted. **c,** Same representation of 14-ADEP bound BsClpP at pH 4.2. The conformation is now compact, and side exit pores (black arrows) start to appear. **d,** Same representation of 5ADEP at pH 5.6. The conformation of BsClpP is compact, and the catalytic triads at the upper and lower heptameric rings become closer and exhibit an inactive configuration. **e,** Same representation of 2ADEP at pH 4.2. The conformation of BsClpP is compressed, and the catalytic triads at the upper and lower heptameric rings are very close. **f,** Same representation of compressed apo BsClpP. The exit pores are wide open for product release. In all panels, the residues maintaining the extended conformation of ClpP are shown as a stick model and labeled. For clarity, the residues in the neighboring subunit of the different heptameric rings are marked with prime (‘) after the residue number, and those in the neighboring subunit of the same heptameric ring are marked with prime (‘) before a single letter code used to represent the amino acid. Positively and negatively charged surfaces are colored blue and red, respectively. The inner surface of the proteolytic chamber is virtually negatively charged (red color), and the acidic proteolytic products must be energetically very unfavorable.

### Comparison with ClpXP structures

Recently, a long-awaited structure of the symmetry-mismatched ATP-dependent Clp protease has been revealed by several independent groups^18–21,26^. The complexes are between a hexameric Clp-ATPase, ClpX or ClpA, and heptameric ClpP, and most of the structures show that six IGF/L-loops of Clp-ATPase bind to six out of seven binding pockets of ClpP with a 10~16° tilt angle of each molecular axis. When we compared the ClpP structures of the ClpXP complex with those of our BsClpP structures, the cryo-EM structures of the extended ClpP at pH 6.5 were similar. More specifically, the monomeric subunit is very similar, while in the oligomeric state of ClpP, the size of the entrance pore is larger in the ADEP complex than in the ClpX complex (Extended Data Fig. 7), showing that ClpX does not induce pore widening of ClpP^19^. The ADEP-binding site overlaps IGF/L-loop binding with an empty site, as shown in ClpXP and ClpAP complexes^18–21^; however, it has been reported that when only 5 IGF/L loops are involved in the interaction with ClpP, the Clp-ATPase conformation has a higher tilting angle (Extended Data Fig. 8)^21^, and this so-called ‘disengaged conformation’ is important for the reaction cycle of the Clp-ATPase. Therefore, we compared our 5ADEP structure with the disengaged conformation. As shown in Extended Data Fig. 8c, the two empty sites in the disengaged conformation of ClpAP are consecutive, whereas the two empty sites in 5ADEP are not (Fig. 2b). Considering the differences in activator binding, the movement and diameter of the entrance pore, and the fact that ADEP cannot actively unfold a protein, the substrate feeding processes realized by two different activators must be different.

## Discussion

The major species of the ATP-dependent complex formed between the Clp-ATPases and ClpP has been reported to be a 2:1 complex^40^, and thus, how the proteolytic products are released from the proteolytic chamber remained mysterious. The pioneering, elegant NMR study coined the concept of the lateral exit pore^29^, and subsequent reports established that the compressed structure of ClpP is the state for product release^13,27,28^ and proposed several factors for structural transitions from the extended state to the compressed state via the compact state^23–26^. Since the first extended structure of ClpP was reported^7^, numerous crystal structures of ClpP from different species and even in different conformational states have been reported ^6,8–13,27^. When we analyzed the crystallization conditions of the reported structures of ClpP, compressed or compact structures were not always but mainly reported for crystals grown under low-pH conditions^13,27^. Therefore, we speculated that there must be a relationship between the physiological conditions and the low pH crystallization that yields compressed structures. ClpP shows a cylindrical shape with doubly capped Clp-ATPases. Thus, its proteolytic chamber is almost a closed compartment, and the proteolytic products must accumulate in it. Therefore, the physicochemical properties of this compartmental environment can be maintained locally. As we described earlier, the peptide products possess millions of new amino- and carboxy-termini that govern the overall pH of this chamber. The average pK_a_ value of all amino acid residues is below pH 6.0, and thus, although it depends on the pI value of the substrates and the composition of amino acids, the local pH value must be approximately pH 6.0. Due to the technical difficulty of direct measuring the pH value of the proteolytic chamber of ClpP during protein hydrolysis, we set up a test experiment on whether substrate hydrolysis indeed drops the pH of the system (Fig. 1). Instead of the compartmentalizing chamber, we used minimal buffering conditions and used a high concentration of the substrate to make a large amount of peptide products. Two model proteins, α-casein and BSA, were degraded by proteinase K, and the pH of the system dropped gradually (Fig. 1). Likewise, when the partially unfolded α-casein was degraded by BsClpP in the presence of ADEP1 (Extended Data Fig. 2b), the pH of the system dropped slowly as the hydrolyzed products were released. We speculate that the pH drop of the proteolytic chamber must occur faster because the volume of the self-compartmentalizing chamber is limited. Therefore, we propose that the main driving force for opening the side exit pore of the Clp-ATPase/ClpP complex is the concentration of protons that regulates the conformational change. The pH change controls countless biological processes, including ATP synthesis, virus maturation, oligomerization, autophagy, and lysosomal activity^41–47^. Here, we add another example, in which the pH-dependent conformational change controls the proteolytic activity of the self-compartmentalizing protease.

There are unique features of ClpP activation by the small ADEP molecules, which are distinguished from those of Clp-ATPases. The ADEP-binding sites share the IGF/L binding region; however, the numbers and positions are different. Previously, only structural information about all 14 ADEP binding or no binding structures has been reported ^11,14,22,27^, and now, we present two other states, 5ADEP (10 ADEPs for a BsClpP tetradecamer) and 2ADEP (4 ADEPs for a BsClpP tetradecamer), which might be energetically stable intermediates among heterogeneous binding states. However, as shown in recent ClpAP cryo-EM structures^21^, depending on the conformational states of ClpA, 6 ADEP-binding sites in the heptameric ClpP ring are occupied in the engaged conformation, and 5 ADEP-binding sites are occupied in the disengaged conformation. The other difference is the entrance pore formed by the N-terminal region of ClpP. Pore widening in the ClpXP complex is not necessary because the ATPase fully unfolds the substrate and translocates it through the narrow pore, whereas ADEP triggers pore opening for substrate translocation^19^. It is known that stably folded model proteins are not degraded by the ADEP-ClpP complex, but partially unfolded substrates such as casein and peptide substrates are well degraded^11,16^. The antibiotic mechanism of ADEP has been established, and the nascent polypeptide chains emerging from the ribosome and, due to intrinsic instability, also the cell division protein FtsZ are the target substrates of the ADEP-ClpP complex^48^. Intriguingly, the intermediate structures 5ADEP and 2ADEP show asymmetric pore shapes due to the molecular asymmetry of ClpP (Fig. 2). In addition, the ClpXP and ClpAP are symmetry-mismatched complexes and thus, the ClpP becomes asymmetric during the reaction cycle. Although the substrate feeding steps realized by ADEP and Clp-ATPases are different, the remaining steps must be similar. Once the substrate reaches the proteolytic chamber, hydrolyzed products are produced, and subsequently, the local pH of the chamber must get lower in both cases. Indeed, negative charges are distributed at the interior of the proteolytic chamber, and the acidic products must be even worse in energetics. Except for the extended state, the strictly conserved catalytic triad showed a distorted configuration, most likely due to the protonation state of His122. The widening of side pores was observed gradually from the extended, fully occupied ADEP-ClpP structure through 5ADEP and 2ADEP to the compressed apo-ClpP structure. In combination with current results and previous reports, we proposed a model for product release (Fig. 6). The product release mechanism of ClpP, initiated by substrate hydrolysis followed by lowering pH, must be similar to both activators, small chemical compounds and Clp-ATPases. However, the enlarged entrance pores of ClpP induced by ADEP binding might be an additional exit site whereas ClpP engaged with processive Clp-ATpase utilizes only side exit pores. Notwithstanding the differences, our current study suggests that the driving force for opening the site exit pore of ClpP is the pH drops coupled with protein hydrolysis. Furthermore, it can be a general concept for self-compartmentalizing proteases.

**Fig. 6:**
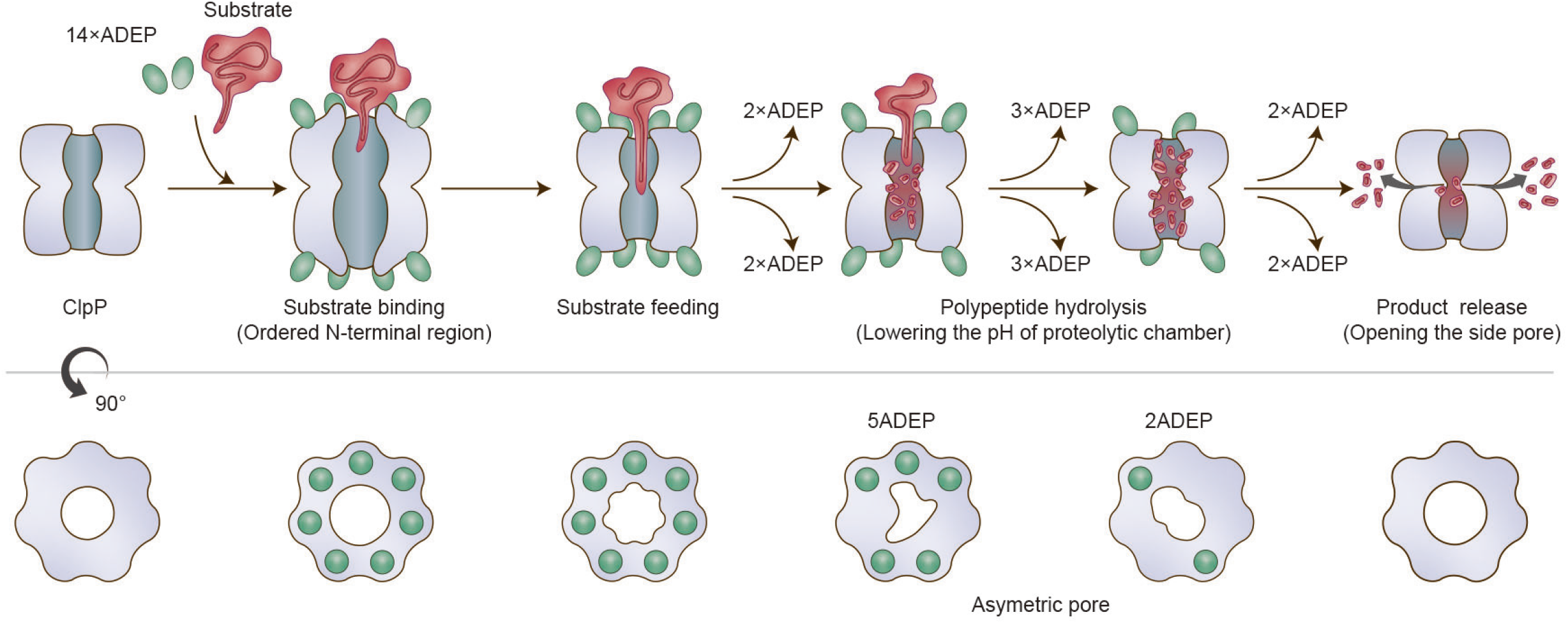
Model proposed for product release of ClpP. Before and immediately after substrate binding, ClpP is in an active extended conformation, and ADEP widens the entrance pore for better substrate feeding. Our previous crystal structure of the 14 ADEP complex showed a disordered N-terminal region^11^ and our current cryo-EM structure of the same complex demonstrates an ordered N-terminal region. In both cases, the entrance pore is enlarged. From the extended structure to the compressed structure, intermediate conformations were observed. Based on the solution cryo-EM analysis, these intermediate conformations represented heterogeneous complex species of BsClpP with different numbers of ADEP molecules bound. Two major species among the intermediate states, 5ADEP and 2ADEP, were crystallized and showed dimensions similar to those of previously known ‘compact’ and ‘compressed’ structures. Both possess asymmetric entrance pores, especially 5ADEP, which is more distorted. 5ADEP is less compressed, and thus, the number and size of the side pores are fewer and smaller than those of a fully compressed structure, respectively. The apo BsClpP at pH 4.2 exhibits compression and larger side exit pores. In the model, the red color of the proteolytic chamber represents the lower pH condition derived from substrate hydrolysis.

## Methods

### Sample preparation

BsClpP protein was purified as previously described^11,27^. For cryo-EM experiments, we further purified the proteins using a Superose™ 6 Increase 10/300 GL (GE Healthcare, 29-0915-96) size-exclusion column pre-equilibrated with 50 mM sodium acetate pH 4.2, 500 mM KCl, and 5% (w/v) glycerol [or 50 mM N-(2-acetamido)iminodiacetic acid pH 6.5, 200 mM NaCl, and 5% (w/v) glycerol]. SDS-PAGE and negative stain electron microscopy were used to assess protein purity and quality. All biological materials are available upon request.

### pH measurement during protein hydrolysis

For *in vitro* pH change measurement, bovine milk α-casein (Sigma, C6780) and bovine serum albumin (Sigma, A7030) were used as substrates. Reactions were carried out with either 100 μg/ml proteinase K (Sigma, P2308) in diluted PBS (1.37 mM NaCl, 0.27 mM KCl, 0.1 mM Na_2_HPO_4_, and 0.18 mM KH_2_PO_4_) containing 1 mM CaCl_2_ or with 0.4 μg/ml BsClpP_14_ in the presence of 11 μM ADEP1 (Cayman Chemical, A-54556A, ~2-fold molar ratio)^11^ in the same PBS at 40 °C. The final concentration of substrate was 80–100 μM, and the final assay volume was 20 ml. The pH was measured every 30 sec for 10 min, every 1 min for 20 min, every 5 min for 210 min, and every 10 min for 60 min, reaching a total of 300 minutes, using a pH meter (Thermo Scientific, 13-644-928).

### Crystallography

A cocrystallization method was used to generate crystals of BsClpP-ADEP complexes by mixing BsClpP and ADEP2 at a 1:1 molar ratio. Crystals of the compressed and compact form of the BsClpP-ADEP complex were grown in hanging drops at 22 °C using 100 mM sodium acetate pH 4.6, 500 mM potassium thiocyanate (for the compressed state), 100 mM sodium citrate pH 5.6, 100 mM Li_2_SO_4_, and 9~11% (w/v) polyethylene glycol 6,000 (for the compact state). Diffraction data were processed with the program HKL2000 (ref. ^49^). The phases of 2ADEP and 5ADEP structures were obtained by molecular replacement using a previously solved compressed BsClpP structure^27^ as a search model with the program MOLREP in the CCP4 suite^50^. Model building and refinement were performed using the programs COOT^51^ and PHENIX^52^, respectively.

### Cryo-EM data collection and processing

The fraction eluted from size-exclusion chromatography was isolated and incubated with 100 μM ADEP1 (Cayman Chemical, A-54556A). In particular, the BsClpP pH 6.5 sample was also treated with 4 mg ml^-1^ amphipol. After 10 min of incubation, a 5 μl drop was applied to a graphene-treated and glow-discharged holey carbon grid (R 1.2/1.3 Quantifoil) for the BsClpP pH 4.2 sample and negatively glow-discharged holey carbon grid (R 1.2/1.3 Quantifoil) for the BsClpP pH 6.5 sample. They were then blotted for 3 sec at 4 °C and 100% humidity with Whatman no. 595 filter paper before being plunge frozen in liquid ethane using a Vitrobot Mark IV system (Thermo Fisher Scientific Inc., USA). Cryo-EM images of frozen-hydrated BsClpP particles were collected at Korea Basic Science Institute with a Titan Krios TEM system (Thermo Fisher Scientific Inc., USA) operated at 300 keV using electron counting mode and automatic data acquisition software (EPU, Thermo Fisher Scientific Inc., USA). Detailed data acquisition conditions and parameters are given in **Supplementary Table 2**.

Cryo-EM images for BsClpP pH 4.2 were processed using RELION 3.0 (ref. ^53^) and CryoSPARC v2 (ref. ^54^). Beam-induced motion correction and dose weighting were performed using MotionCor2 v1.2.1 (ref. ^55^), and CTF estimation was performed using Gctf v1.06 (ref. ^56^). Then, 341,414 particle images were selected from 964 micrographs after reference-free 2D class averages. The 3D initial model was generated by CryoSPARC-implemented Ab initio reconstruction. Multiple rounds of successive 3D classifications were performed, and 179,322 homogeneous particles were selected for further processing. Homogeneous refinement with D7 symmetry then yielded a consensus map at 3.6 Å. Of note, particles displayed a significant preferred orientation, showing a dominant top view. Selected particles with angular information were transferred to RELION for further 3D classification without an angular orientation search. To evaluate the possible heterogeneity of ADEP binding, we performed focused 3D classification around ADEP binding sites with expanded particles with C7 symmetry. We only observed the all-ADEP and apo states from the analysis, and thus, we clustered particles into apo or all-ADEP states. Finally, each selected particle set was imported to CryoSPARC and refined at 3.4 Å.

The images of BsClpP pH 6.5 were processed using CryoSPARC v2 (ref. ^54^). We originally analyzed ADEP binding heterogeneity in the dataset of a 1:3 mixture of BsClpP and ADEP at pH 6.5 using the same procedure as described above in BsClpP pH 4.2. However, particles were homogeneous, as all ADEP binding sites were occupied in the dataset. Therefore, we obtained two separate imaging sessions for apo (1,130 movies) and ADEP-bound structures (1,182 movies) at pH 6.5. The recorded movies were subjected to motion correction and CTF estimation using patch-motion correction and Gctf v1.06 in CryoSPARC. For the apo map at pH 6.5, a total of 108,389 particle images were selected after 2D class averages. After subsequent heterogeneous refinement for suitable particles, 78,413 particles were finally used for 3D reconstruction at 3.2 Å resolution. For the ADEP bound map at pH 6.5, a total of 153,078 particle images were selected after 2D class averages. After subsequent heterogeneous refinement, 138,976 particles yielded the map at 3.1 Å resolution. All cryo-EM images were processed using computing resources at the Center for Macromolecular and Cell Imaging, Seoul National University.

### Molecular modeling

Both BsClpP pH 4.2 and pH 6.5 structural models were built manually in COOT^51^ by referring to the BsClpP crystallography structure (PDB ID: 3TT6 for the pH 4.2 model and 3KTJ for the pH 6.5 model) and refined using phenix.real_space_refine in the PHENIX software suite^57^. Additionally, the quality of the final models was evaluated using the comprehensive model validation section and MolProbity in PHENIX.

## Acknowledgments

We thank the staff at beamlines 5C and 11C at the Pohang Accelerator Laboratory in South Korea and beamline NW12 at the Photon Factory in Japan for their help with X-ray data collection. We also thank the staff of the cryo-TEM facility at the Korea Basic Science Institute and the Center for Macromolecular and Cell Imaging, Seoul National University. This study was supported by National Research Foundation of Korea (NRF) grants from the Korean government (grant Nos. 2020R1A2C3008285, 2020R1A5A1019023, and 2021M3A9I4030068 for HKS; 2021M3A9I4021220, 2019M3E5D6063871, and 2020R1A5A1018081 for SHR) and the Deutsche Forschungsgemeinschaft (German Research Foundation, DFG) TRR 261, project-ID 398967434.

## Author contributions

L. K. and B.-G.L. performed sample preparation for biochemical experiments; B.-G.L. and M. K.K. prepared the crystals and solved the X-ray structures; L.K., M.K.K, D.H.K. and S.-H.R. performed the cryo-EM studies; L.K., B.-G.L., H.K., H.B.-O., S.-H.R., and H.K.S. analyzed the data; H.B.-O. discussed results and edited the manuscript; and L.K., B.-G.L., and H.K.S. designed the experiments and wrote the manuscript.

## Competing interests

The authors declare no competing interests.

## Data availability

The cryo-EM maps have been deposited in the EMDB: ADEP1-BsClpP complex at pH 4.2 (EMD-31561), apo-BsClpP at pH 4.2 (EMD-31562), ADEP1-BsClpP complex at pH 6.5 (EMD-31559), and apo-BsClpP at pH 6.5 (EMD-31560). The coordinates have been deposited in the PDB: crystal structures – 2ADEP (7P80) and 5ADEP (7P81); cryo-EM structures – ADEP1-BsClpP complex at pH 4.2 (7FER), apo-BsClpP at pH 4.2 (7FES), ADEP1-BsClpP complex at pH 6.5 (7FEP), and apo-BsClpP at pH 6.5 (7FEQ).

## Code availability

Not applicable.

